# Upregulation of DEK Expression in Uterine Myomas and Cervical Cancer as a Potential Prognostic Factor

**DOI:** 10.1101/2024.08.12.607532

**Authors:** Amelia Janiak, PengCheng Tan, Ferdinand Kappes, Felice Petraglia, Chiara Donati, Xinyue Liu, Renata Koviazina, Fangrong Shen, Anastasia Tsigkou

**Author notes:** **Corresponding Author:** Anastasia Tsigkou, **Email:**. **Co-corresponding Author:** Fangrong Sheng **Email:**.

## Abstract

The aim of this pilot study is to investigate the role of the DEK protein as a potential prognostic marker in gynecological tumors, specifically focusing on uterine myomas and cervical cancer. The study cohort comprised Chinese female patients manifesting with menorrhagia and pelvisalgia, from whom neoplastic and adjacent non-neoplastic tissue specimens were procured during surgical intervention for either leiomyomas or cervical carcinoma. DEK protein and messenger RNA (mRNA) levels were measured across normal uterine tissue, uterine myomas, and cervical cancer tissues using Western blotting, immunohistochemistry, and quantitative real time polymerase chain reaction (qRT-PCR). Results revealed a marked increase in DEK protein expression in cervical cancer tissues, moderate expression in uterine myomas, and minimal levels in normal uterine tissues. Statistical analyses confirmed significant differences in DEK protein expression between tissue types, though mRNA expression differences did not reach statistical significance. These findings suggest DEK’s involvement in tumor development and suppression, making it a promising biomarker for early detection in gynecological tumors. Further research is needed to elucidate DEK’s mechanisms in gynecological tumorigenesis and its potential as an early biomarker, addressing critical need in women’s health.

## 1. Introduction

Tumors of the female reproductive system are very common, including uterine myomas as the most frequent benign tumor, while cervical cancer the most common malignancy (1, 2). Uterine myomas, also known as fibroids, arise from the overgrowth of smooth muscle layer and connective tissue in the uterus and can be found in about 50% of women in reproductive age (1, 3). Although generally non-malignant, such conditions can manifest with clinical complications, including pelvisalgia, menorrhagia, and fertility-related challenges (4, 5). Conversely, cervical carcinoma arises from the cellular architecture of the cervix and represents a significant health issue in the absence of timely diagnosis and intervention (6).

Current screening methods for cervical cancer and fibroids utilize various diagnostic tools, including transvaginal ultrasound, pelvic examinations, magnetic resonance imaging (MRI), and Pap smears. For cervical cancer, Pap smears are the primary screening test recommended every three years for women aged 21 to 65 (7). For women over 30, HPV testing—particularly for high-risk types like HPV-16 and HPV-18—is often combined with Pap smears because these high-risk HPV types are strongly associated with the development of cervical cancer (7). However, these traditional methods are not without limitations. Pap smears have a sensitivity of only about 50%, with estimates ranging from 30% to 87% (8). This variability can lead to false positives and negatives, resulting in gaps in detection and follow-up care. In fact, a meta-analysis by Spence et al. showed a false negative rate of cytologic testing of 35.5% on average (9). While HPV testing improves sensitivity to up to 96% and identifies high-risk strains before cytological abnormalities appear (10), its roughly 90% specificity means that it still falls short of fully addressing the challenges of comprehensive early detection as it may still miss some precancerous lesions or fail to identify all high-risk individuals.

For uterine myomas, transvaginal ultrasound and MRI are considered the optimal methods to assess uterine myomas number, volume, echostructure, location, relation with endometrial cavity and uterine layers, vascularization, and differential diagnosis with other benign and malignant myometrial pathologies (11). Transvaginal ultrasound is widely used due to its accessibility and real-time imaging capabilities; however, its effectiveness can be limited by operator dependency and may be compromised in patients with obesity or large fibroids that obscure the view (12). MRI offers superior resolution and a comprehensive view of the uterus, but it is more costly, less accessible, and has limitations such as longer scan times and the need for patient cooperation (12). Additionally, MRI may not always accurately distinguish between myomas and other similar pathologies, which can lead to misdiagnosis (13).

To enhance diagnostic accuracy, integrating additional biomarkers into existing screening protocols has shown promise. For instance, p16/Ki-67 dual staining has proven valuable in improving the reliability of cervical cancer screening, indicating that incorporating protein and gene markers could yield a more nuanced and personalized approach (14). Similarly, in the context of uterine fibroids, genetic testing can provide insights into individual risk and potential progression by detecting specific gene mutations, such as MED12, HMGA2, and FH (15). However, the addition of more genetic and protein markers is needed to further improve diagnostic precision and patient outcomes.

Given an urgent need for improved screening methods and the rising prevalence of gynecological tumors with profound impact on quality of life, including physical, emotional, and psychological well-being (16, 17), current research is increasingly focused on novel agents that target the DNA damage response (DDR). DDR comprises a set of signaling pathways for the detection and repair of DNA damage, including machinery mediating DNA repair, cell cycle regulation, replication stress responses and apoptosis (18). Indeed, the progression of cervical cancer is associated with an increased genetic instability, which is primarily caused by the DNA damage and breakage (19). Enhanced comprehension of DDR processes offers potential for cancer treatment by elucidating cellular and molecular signaling mediators pivotal in tumor development and progression.

One such promising marker within this context is the rather under-investigated, multifunctional chromatin-associated oncogene DEK. The DEK gene is located in the chromosome 6p22.3 band and encodes for a 43 kDa, highly conserved nuclear protein predominantly expressed in malignant and actively dividing cells (20, 21). Initially recognized for its role in chromatin organization and gene regulation, DEK has been implicated in various cellular processes such as DNA damage repair, RNA transcriptional regulation, mRNA splicing, and DNA replication (22, 23, 24, 25, 26, 27). Recent studies have also demonstrated that elevated DEK levels promote proliferation, motility, invasion, and tumorigenesis (26, 28, 29), prompting investigations into various cancer types, which showed that DEK is upregulated in acute myeloid leukemia, retinoblastoma, glioblastoma, melanoma, and in a growing number of other tumor types (30, 31). In addition, suppressing DEK and nuclear factor kappa B has been shown to have the potential to arrest the cell cycle in the G0/G1 phase, resulting in fewer cells in the G2/M phase, and increased apoptosis and cell senescence in CaSki cervical cancer cells, suggesting that DEK may have an oncogenic function in the development and progression of tumors (32). Similar conclusions were drawn by Xu et al., who confirmed DEK as an oncoprotein in cervical cancer, correlating with FIGO staging and tumor type, and demonstrated that silencing DEK inhibited cancer cell proliferation, migration, and invasion by downregulating Wnt/β-catenin and MMP-9, increasing GSK-3β activity, and impairing tumorigenicity in a mouse xenograft model (33).

The oncogenic role of DEK is part of a broader mechanism where the buildup of genetic damage, including the activation of proto-oncogenes and the inactivation of tumor-suppressor genes, propels the transformation of healthy cells into malignant ones (34). Proto-oncogenes, such as DEK, which regulate cell differentiation and proliferation, can cause neoplastic transformation if mutated, a process known as activation (35). Activation occurs through various genetic pathways such as transduction, insertional mutagenesis, amplification, point mutations, and chromosomal translocations, resulting in deregulated proto-oncogenes that provide a proliferative benefit to the cell (35). As on oncogene, DEK has been shown to promote tumorigenesis by interfering with cell division, affecting DNA repair, inhibiting cell differentiation, senescence, apoptosis, and cooperating with other oncogenes (22, 36). Supporting this, studies have demonstrated DEK’s significant role in tumorigenesis. Han et al. discovered that DEK is significantly involved in the proliferation of serous ovarian cancer cells, with high DEK expression levels correlating with an increased Ki-67 proliferation index (37), while Privette et al. demonstrated that DEK oncogene stimulates cellular proliferation via Wnt signaling in Ron receptor-positive breast cancers (38).

While DEK overexpression is frequently linked to malignant tumors, it is also observed in various benign tumors. For instance, studies with mouse models have shown that DEK knockout mice are partially resistant to the formation of benign skin papillomas, indicating DEK’s role in early tumorigenesis and involvement in suppressing tumor growth (39). However, other studies, such as Riveiro-Falkenbach et al., have found negligible DEK expression in benign lesions, presenting conflicting evidence about DEK’s role in non-malignant neoplasms (40). Additionally, research on DEK expression in melanocytic lesions found low or no DEK expression in most benign nevi and melanoma in situ (41). Despite these mixed findings, there is still limited information on DEK in benign tumors, and further research could provide valuable insights into its role in these tumors.

Given DEK’s critical role in tumorigenesis and DNA repair, along with the limited research on its function in benign tumors and the need for more accurate diagnostic biomarkers for Pap smears, our study aimed to evaluate DEK expression as a potential prognostic factor in both benign gynecological tumors and malignant gynecological cancers. Through immunohistochemical, Western blot, and quantitative real time polymerase chain reaction (qRT-PCR) analyses, we observed significant DEK upregulation in cervical carcinomas, moderate expression in uterine myomas, and negligible expression in normal uterine tissues, with statistically significant differences across these tissues.

## 2. Materials and Methods

### 2.1 Tissue Samples

This study aims to explore differences in DEK expression between benign uterine myomas and cancer tissues. Due to limitations in patient recruitment, this work serves as a pilot study. Specimens of myoma and myometrium were collected according to the guidelines of the institutional review board of No. 1 Affiliated Hospital Suzhou University, department of Obstetrics and Gynecology, China and Duke Kunshan University, Kunshan, China. Informed written consent was obtained from all patients prior to surgery. Tissue samples were collected during surgery between January and June 2023 from ten women with uterine myomas, four with cervical cancer, and three with normal uterine tissue; each sample came from a different patient. Surgeries were conducted due to women experiencing heavy bleeding, abdominal pain, and dysmenorrhea during their luteal phase. An ultrasound and pap smears were performed for diagnostic purposes. Uterine myoma and cancer samples were not matched with control samples. The normal tissues were obtained from patients with no prior history of fibroids, cancer, or endometriosis. For each patient, uterine leiomyoma risk factors - such as age, number of leiomyomas, VAS index, and the menstrual cycle phase (middle to late proliferative) - were recorded. None of the patients had received steroid hormone therapy (OAC, IUD, or HRT) for at least 3 months prior to surgery. Endometrial tissue (0.4–1.2 g) for uterine myomas and normal tissues was scraped from the fundus uteri using a surgical knife immediately after surgery. Cone biopsy was obtained from cervical cancer samples. The samples were snap-frozen in liquid nitrogen and stored at −80°C. When a leiomyoma was present, tissue was collected from the opposite side of the uterus. Pathological examination at the No. 1 Affiliated Hospital confirmed the presence of uterine myomas after myomectomy and assessed the pathological stage and histological subtype for fibroid and cancer samples according to the 1988 FIGO criteria, with all fibroids staged at FIGO I-III and all cancers staged at FIGO I-III (43, 52). Due to the small sample size, no a priori power analysis was performed, and all available samples were analyzed.

### 2.2 Immunohistochemical Analysis

#### 2.2.1 Immunohistochemical Studies of Tissue Explants

Immunohistochemical staining for DEK (1:1000, DEK (Abcam, ab26) proteins was performed using formalin-fixed and paraffin-embedded tissue slides, a detection kit (DAKO ChemMate, DAKO, Glostrup, Denmark) and a semi-automated stainer (DAKO TechMate, DAKO, Glostrup, Denmark) according to the specifications of the manufacturer. For antigen retrieval the slides were treated in a PT Link module (DAKO, Glostrup, Denmark) using the EnVision™ FLEX Target Retrieval Solution, Low pH (DAKO, Glostrup, Denmark). Quantification of DEK expression was performed using QuPath image analysis software, which facilitated semi-automated analysis of DAB-positive cells. Two independent observers evaluated the samples, and the optical staining intensity was determined (graded as 0: no staining; 1: weak; 2: moderate; and 3: strong staining) along with the percentage of positive cells (0: no staining; 1: <10%; 2: 11-50%; 3: 51-80%; 4: >81% of cells). The IRS score was calculated by multiplying these two values.

### 2.3 Western Blot Analysis

#### 2.3.1 Protein Extraction

Tissues were dissected on ice and transferred to microcentrifuge tubes. For 100 mg of tissue, 400 µL of ice-cold RIPA lysis buffer with PMSF (Beyotime, P0013B) was added, and the tissue was homogenized using a pestle and a mortar. Additional 600 µL of lysis buffer was added during the homogenization process. The sample was kept on ice and agitated on an orbital shaker for 1.5 hours. Subsequently, the tubes were centrifuged at 4°C. The supernatant containing extracted proteins was collected into a fresh tube and stored at −80°C.

#### 2.3.2 Western Blot

Total protein concentration was determined using the Enhanced BCA Protein Assay Kit (Beyotime, China, P0009). The proteins and deionized water were mixed to make sure each sample had the same concentration with a total of 20 μL or 30 μL. 5 μL of 5x SDS-PAGE Loading Buffer (NCM Biotech,WB2001) was then added to each sample, followed by vortex, incubation at 95°C for 5 min and centrifugation at 1,000g for 10 seconds. Total protein (16 μg per lane) was separated using a SDS-polyacrylamide gel and transferred onto a nitrocellulose membrane. SeeBlue Plus2 Pre-Stained Standard (Invitrogen, Karlsruhe, Germany) was used as a marker. The membranes were incubated with primary anti-DEK (rabbit, 1:2000; Abcam, ab26) and anti-GAPDH (rabbit, 1:1000; Beyotime, AG0122) followed by incubation with the corresponding secondary antibodies (HRP-conjugated goat anti-rabbit IgG, 1:5000, MULTISCIENCE, GAR0072). The bands were visualized by incubating the membrane with ECL solution (Tanon^TM^ High-sig ECL Western Blotting Substrate (Biotanon,180-501) and examined in a pre-cooled chemiluminescence imaging system (Bio-Rad Laboratories, CA, USA) according to the manufacturer’s instructions. Image analysis was performed with the ImageJ to obtain quantifiable DEK protein expression levels, normalized to the expression of the reference gene (GAPDH).

### 2.4 qRT-PCR Analysis

#### 2.4.1 RNA Extraction

Tissue samples stored at −80°C were crushed until a fine powder using a mortar and a pestle. Approximately 100 mg powder was recovered from each sample and placed into a new cold microtube. Then, RNA was extracted using the TRIzol reagent (ThermoFisher Scientific, 15596026) protocol provided by the manufacturer, using TRIzol reagent, chloroform (Titan,G75915B) and isopropanol (Titan, G75885B). After RNA extraction, 20 μL of DEPC water (DNase, RNase free) (damas life, G8010-500ml) was added to the sample and it was stored at −80°C.

#### 2.4.2 cDNA Synthesis

cDNA synthesis was carried out using HiScript II 1st strand cDNA synthesis kit (Vazyme, R211-02). About 500 ng of total RNA were reverse transcribed with 200 U/μl of M-MLV reverse transcriptase (Invitrogen), RNase Out, 150 ng random hexamers and 10 mM dNTPs according to the manufacturer’s instructions. RNA was denatured at 65°C for 5 min and subsequently kept on ice for 1 min. After adding the enzyme to the RNA primer mixes, samples were incubated for 10 min at 25°C to allow annealing of the random hexamers. Reverse transcription was performed at 37°C for 50 min followed by inactivation of the reverse transcriptase at 70°C for 15 min.

#### 2.4.3 qRT-PCR

qRT-PCR was utilized in human tissue samples to analyze gene expression levels of two proteins: DEK and GAPDH. Relative quantification of transcription levels was carried out by real-time PCR analyses using the Applied Biosystems 7300 real-time PCR system (Applied Biosystems) and ChamQ universal SYBR qPCR Master Mix kit (Vazyme,Q711-02) according to the manufacturer’s instructions. Sequences for primers used in the procedure were found in the scientific literature and commercially synthesized by Sangon Biotech. Primer sequences for DEK were: 1) Forward: 5’-TGGGTCAGTTCAGTGGCTTTCC-3’, 2) Reverse: 5’-CTCTCCAAATCAAGAACCTCACAG-3’, and for GAPDH were: 1) Forward: 5’-ACAACTTTGGTATCGTGGAAGG-3’, 2) Reverse: 5’-GCCATCACGCCACAGTTTC-3’. The amplification reaction mixture (total volume of 20 μL) contained 2 × ChamQ Universal SYBR qPCR Master Mix, 0.4 μL of forward primer (10 μΜ), 0.4 μL of reverse primer (10 μΜ), 2 μL of template cDNA, and DEPC-H2O (added to the final volume). Incubation conditions were: 1) initial denaturation: 95°C/30 sec; 2) cycling reaction: 40 cycles of 95°C/10 sec and 60°C/60 sec; 3) melting curve: 95°C/15 sec, 60°C/60 sec, 95°C/15 sec.

### 2.5 Statistical Analysis

To determine the statistical significance of DEK protein expression differences among the three tissue types, two statistical tests were utilized to analyze Western blot results. First, one-way ANOVA test was performed to determine if there were any overall significant differences in DEK expression among the groups. Following a significant ANOVA result, post-hoc Tukey’s Honest Significant Difference (HSD) test was conducted to identify specific pairwise differences between the groups.

The results of the qRT-PCR were expressed as 2^-ΔΔCt, which is the fold change in gene expression relative to the control (normal uterine tissue), calculated as the difference between the delta cycle threshold (ΔCt) of the target gene and the ΔCt of the reference gene (GAPDH). To compare gene expression patterns, relative gene expression was graphed in GraphPad Prism 10. Statistical analysis was conducted using Kruskal-Wallis H test and unpaired Mann-Whitney U test. Kruskal-Wallis H test was employed to assess the presence of overall differences in expression among normal uterine tissue, uterine myoma, and cervical carcinoma samples, while unpaired Mann-Whitney U test was used to further examine and identify specific group differences.

## 3. Results

Cervical intraepithelial neoplasia of different stages was obtained (cervical intermediate differentiated adenocarcinoma IB1 = 2, cervical low-grade squamous cell carcinoma IIIC1 = 1, cervical low-grade squamous cell carcinoma IB (specific IB123 pathology unknown) = 1). The median age at the time of diagnosis for patients with cervical cancer was 44, with 28-53 age range. The clinicopathological features of uterine myomas are summarized in Table I.

**Table 1.**
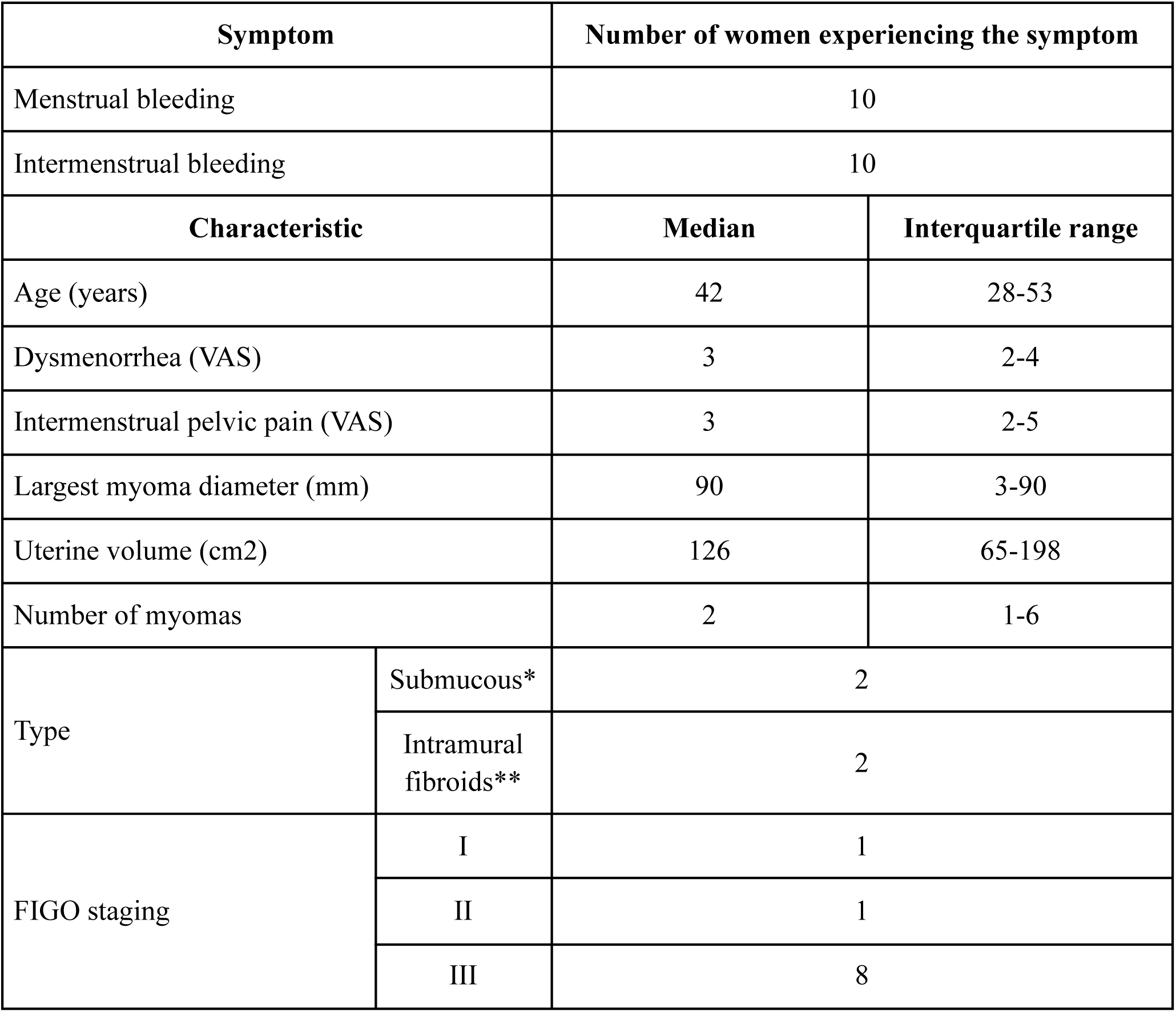
Clinical Characteristics of Women With Uterine Myomas (n=10). VAS stands for visual analog scale. *submucosal fluid with intramural extension smaller or bigger 50%; ** in contact with the endometrium, but not extending into the uterine cavity or serous surface.

| Symptom |  | Number of women experiencing the symptom |  |
| --- | --- | --- | --- |
| Menstrual bleeding |  | 10 |  |
| Intermenstrual bleeding |  | 10 |  |
| Characteristic |  | Median | Interquartile range |
| Age (years) |  | 42 | 28-53 |
| Dysmenorrhea (VAS) |  | 3 | 2-4 |
| Intermenstrual pelvic pain (VAS) |  | 3 | 2-5 |
| Largest myoma diameter (mm) |  | 90 | 3-90 |
| Uterine volume (cm2) |  | 126 | 65-198 |
| Number of myomas |  | 2 | 1-6 |
| Type | Submucous* | 2 |  |
|  | Intramural fibroids** | 2 |  |
| FIGO staging | I | 1 |  |
|  | II | 1 |  |
|  | III | 8 |  |

### 3.1 Immunohistochemical Analysis

Expression levels of DEK protein in normal uterine, uterine myoma and cervical carcinoma tissue samples were determined via immunohistochemical counterstaining. Representative immunohistochemical images are shown in Figure 1A-E and the IRS scores are presented in Figure 1F. The data indicates a significant upregulation in DEK protein expression in cervical cancer tissue, with a high final IRS score of 12, indicating strong positive expression. In uterine myoma tissue, DEK expression is present but at a lower level, with a final IRS score of 2, indicating positive expression. Normal tissue shows little to no DEK expression, with a final IRS score of 0.

**Figure 1.**
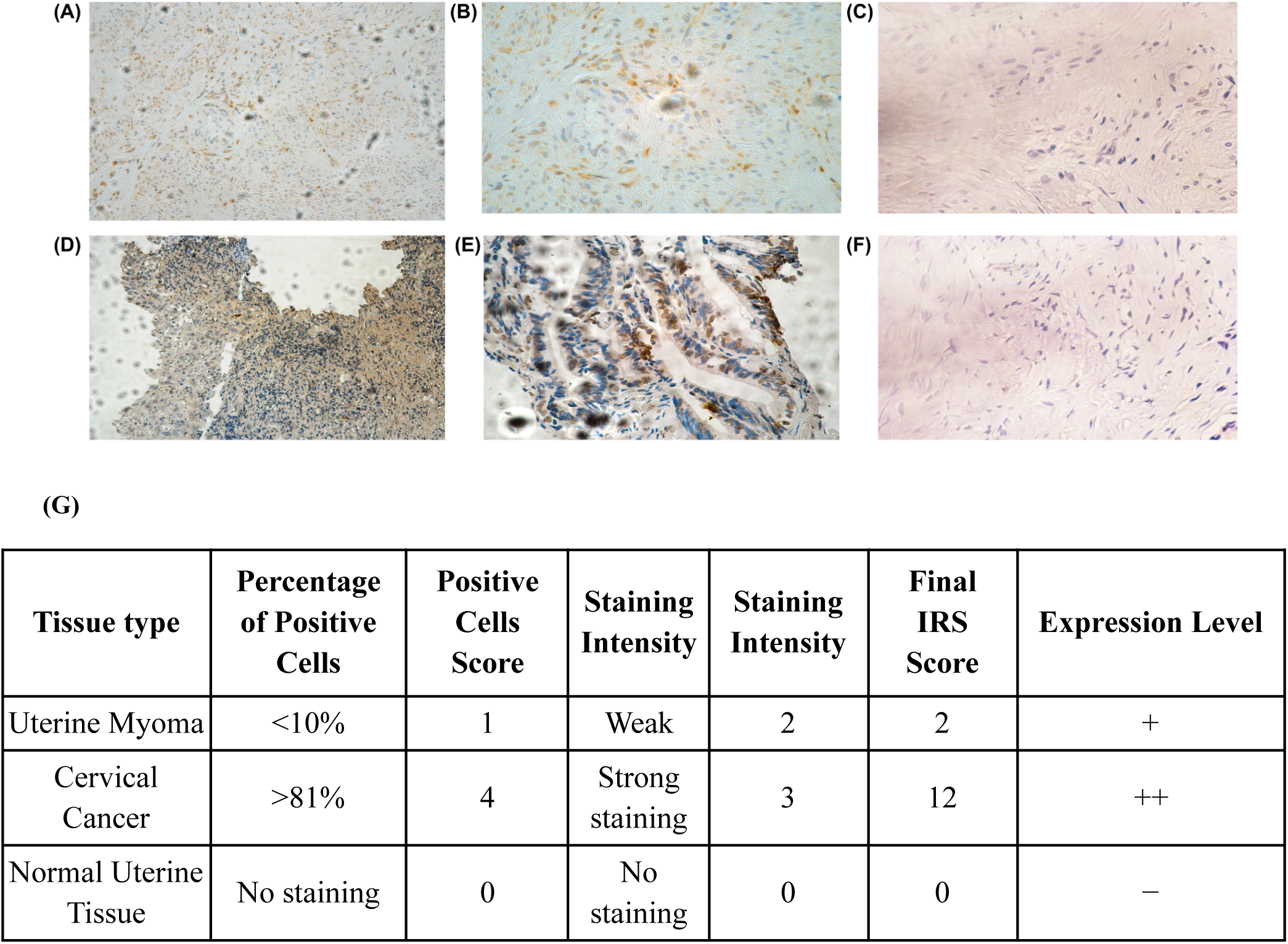
Immunohistochemical Staining and Analysis of DEK Expression in Uterine Myomas and Cervical Cancer Tissues. Immunohistochemical staining for DEK (1 mg/ml, Abcam) was conducted on formalin-fixed, paraffin-embedded tissues using a detection kit (DAKO ChemMate) and a semi-automated stainer (DAKO TechMate) according to the specifications of the manufacturer. (A) Nuclear staining of DEK, uterine myoma (x20). (B) Nuclear staining of DEK, uterine myoma (x40). (C) Negative control for nuclear staining of DEK, uterine myoma (x40). (D) Nuclear staining of DEK, cervical cancer (x20). (E) Nuclear staining of DEK, cervical cancer (x40). (F) Negative control for nuclear staining of DEK, cervical cancer (x40). (G) To quantify DEK expression in uterine myomas and cervical cancer tissues immunohistochemical staining analysis was conducted. The IRS score was calculated by multiplication of the optical staining intensity (0, no staining; 1, weak; 2, moderate; 3, strong staining) and the percentage of the positive stained cells (0, no staining; 1, <10%; 2, 11-50%; 3, 51-80%; 4, >81% of the cells). The expression level of DEK was then categorized into three levels: negative (−), positive (+), and strongly positive (++).

### 3.2 Western Blot Analysis

Western blotting was used to investigate DEK protein expression levels in normal uterine tissue, uterine myoma tissue and cervical cancer tissue samples. The results were analyzed using a computer program (ImageJ) and were normalized to GAPDH expression.

As shown in Figure 2, relative DEK protein expression level is highest in cervical cancer tissue (∼5.7), intermediate in uterine myoma tissue (∼4.0), and lowest in normal uterine tissue (∼1.0). Statistical analysis with ANOVA test showed a significant difference in DEK expression among the tissue types (*p* = 0.0038). Post-hoc analysis with Tukey’s HSD test confirmed significant differences between normal and uterine myoma tissues (*p* < 0.05) and between normal and cervical cancer tissues (*p* < 0.01), but not between uterine myoma and cervical cancer tissues (*p* > 0.23).

**Figure 2.**
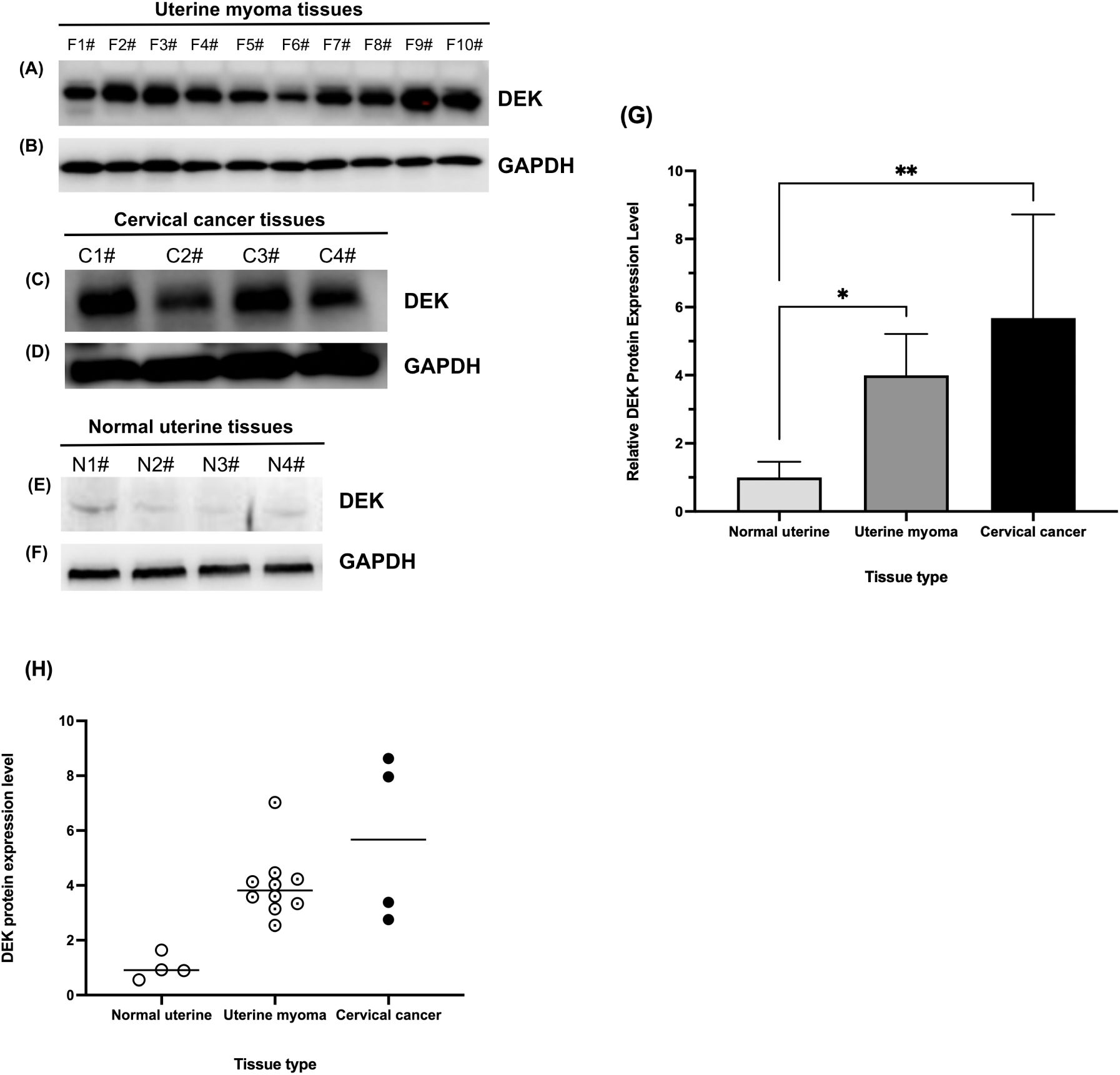
Relative DEK Protein Expression in Normal Uterine Tissue, Uterine Myoma Tissue, and Cervical Cancer Tissue. Western blot analysis of protein expression: (A) DEK in uterine myoma (F1#-F10#), (B) GAPDH in uterine myoma (F1#-F10#), (C) DEK in cervical cancer (C1#-C4#), (D) GAPDH in cervical cancer (C1#-C4#), (E) DEK in normal uterine (N1#-N4#), (F) GAPDH in normal uterine (N1#-N4#). Antibodies: DEK (rabbit, 1:2000, Abcam), GAPDH (rabbit, 1:1000, Beyotime). (G) Quantification of DEK and GAPDH protein levels from the Western blots in panels A-F using ImageJ gel analysis program. Data are presented as mean and standard deviation (error bars). Statistical analysis was performed using one-way ANOVA (*p* = 0.0038) and Tukey’s HSD test for pairwise comparisons: normal vs. uterine myoma (*p* < 0.05), normal vs. cervical cancer (*p* < 0.01), uterine myoma vs. cervical cancer tissue (*p* > 0.23). *, *p* < 0.05; **, *p* < 0.01. (H) Relative DEK protein expression within individual biological samples.

### 3.3 qRT-PCR

To analyze DEK expression at the transcriptional level, mRNA was extracted from samples of cervical cancer, uterine myomas and normal uterine tissue, transcribed into cDNA, and followed by qRT-PCR analysis.

Figure 3 demonstrates that in normal uterine tissue, relative DEK mRNA expression is relatively low, with an average at 1.031. The uterine myoma tissue exhibits higher relative expression, with an average close to 1.085, indicating an upregulation in this tissue type. In cervical cancer tissue, the relative DEK mRNA expression is similar to that of uterine myoma tissue, averaging around 1.091. These results suggest that similarly to protein levels, DEK mRNA is upregulated in uterine myoma tissue and cervical cancer tissues, which show similar expression levels. Statistical analysis, however, demonstrated that there are no statistically significant differences in DEK expression levels among the three tissue types (Kruskal-Wallis H (*p* = 0.503); unpaired Mann-Whitney U test: normal vs. uterine myoma tissues (*p* = 0.349), normal uterine vs. cervical cancer tissues (*p* = 0.571), uterine myoma vs. cervical cancer tissues (*p* = 0.566)).

**Figure 3.**
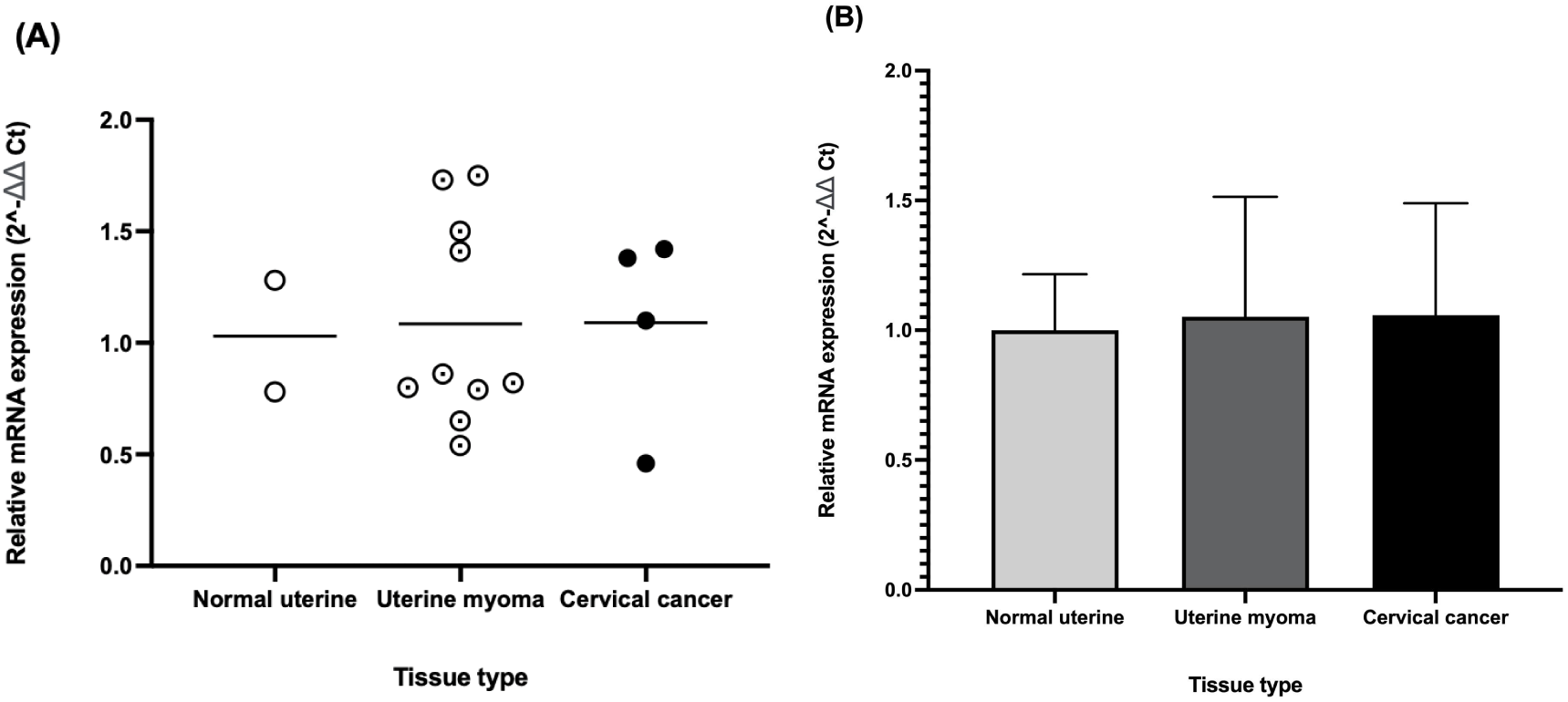
Relative DEK mRNA Expression Levels in Normal Uterine Tissue, Uterine Myoma tissue, and Cervical Cancer Tissue. Relative quantification was performed using the Applied Biosystems 7300 real-time PCR system and ChamQ Universal SYBR qPCR Master Mix. Primers for DEK and GAPDH were sourced from literature and synthesized by Sangon Biotech. (A) Relative DEK mRNA expression (2^-ΔΔCt). (B) Mean relative DEK mRNA expression (2^-ΔΔCt) with standard deviations (error bars). No statistically significant differences in DEK expression levels among the three tissue types were found.

## 4. Discussion

Our pilot study provides evidence of varying DEK expression in normal uterine tissue, uterine myomas, and cervical cancer tissues, highlighting DEK’s potential role in gynecological tumorigenesis and its value as a prognostic marker. Immunohistochemical analysis shows significant upregulation of DEK in cervical cancer tissues, moderate expression in uterine myomas, and negligible expression in normal uterine tissues. Western blot analysis, normalized to GAPDH, corroborates these findings with DEK protein expression highest in cervical cancer tissues, lower in uterine myomas, and lowest in normal tissues. Consistent with protein expression patterns, qRT-PCR analysis indicates elevated DEK mRNA levels in uterine myomas and cervical cancer tissues compared to normal uterine tissues. Statistical analysis of Western blot data, conducted using one-way ANOVA and Tukey’s HSD post-hoc test, revealed significant differences in protein expression across groups. However, no statistically significant differences in DEK mRNA expression were observed, potentially due to the limited sample size typical of a pilot study.

These findings may suggest that DEK plays a distinct role in different gynecological tissues and tumor types. The marked upregulation of DEK in cervical cancer tissues underscores its potential as an oncogene. DEK’s overexpression in cervical cancer cells aligns with previous studies demonstrating its involvement in promoting cell proliferation, motility, invasion, and tumorigenesis (26, 28, 29, 33, 44). In reproductive system cancers, such as breast cancer, this upregulation may enhance cancer proliferation and invasiveness by activating the Wnt signaling pathway through increased levels of active β-catenin and Wnt target genes, including cyclin D1 and c-Myc (45). It can also promote angiogenesis and metastasis by activating the PI3K/AKT/mTOR pathway (56). Taken together, these mechanisms highlight DEK’s critical role in driving cancer progression and underscore its potential as a therapeutic target in gynecological malignancies.

The intermediate levels of DEK expression in uterine myomas suggest that DEK may act as a tumor suppressor in these benign tumors. This elevated DEK expression in myomas may be linked to a compensatory response to control cellular proliferation and maintain tissue integrity, counteracting the potential for malignant transformation. Specifically, DEK has been found to counteract DNA damage that arises from replication stress, such as stalled or collapsed replication forks, which often occur at fragile chromosomal sites (46). This happens in a manner similar to other DNA repair proteins, such as FANCD2 and RAD51, which help stabilize stalled replication forks and prevent double-strand break formation (22, 47). By preventing excessive DNA damage and facilitating proper fork progression, DEK may function as a tumor suppressor and its elevated expression may reflect a compensatory mechanism to maintain genomic stability and prevent the transition from benign to malignant states.

The negligible expression of DEK in normal uterine tissues aligns with its predominant expression in rapidly dividing and cancerous cells, as previously reported (21). The low levels of DEK in normal tissues suggest that its upregulation is associated with pathological conditions, reinforcing the concept that DEK functions as an oncogene or tumor promoter in abnormal cellular environments.

Although our study did not reveal statistically significant differences in DEK gene expression across normal, benign, and malignant gynecological tissues, we did observe significant variations in DEK protein expression patterns. These findings suggest that DEK could be a promising biomarker for early screening, with potential use in Pap smears to detect DNA damage or early tumorigenic changes. By monitoring DEK protein expression levels, it might be possible to detect early genetic instability associated with the onset of fibroids and cervical cancer. This capability opens up new possibilities for early screening methods at the protein level, enabling timely intervention before tumors progress to advanced stages. Specifically, assessing DEK protein expression could help identify women at higher risk for developing these conditions, facilitating early diagnosis and improving patient outcomes. Incorporating DEK as a biomarker in early screening protocols, as suggested by other studies for various cancer types (39, 40, 41), could significantly advance the early detection and management of gynecological tumors, addressing a crucial need in women’s health. However, it is important to note that this is a pilot study with a limited sample size, and future research should expand the sample size to validate our findings and determine whether DEK mRNA expression exhibits significant differences across tissue types, potentially solidifying DEK’s role as a biomarker in gynecological oncology.

These findings also underscore the need for further research to elucidate the precise mechanisms by which DEK contributes to gynecological tumorigenesis. Such research could focus on the interplay between DEK, p53, TGF-β, and S1P with particular attention to how DEK influences p53 and TGF-β receptor stability as well as their downstream signaling (50, 51). Understanding these interactions is important since TGF-β signaling plays a critical role in cancer progression, influencing diverse cellular processes such as cell growth, differentiation, apoptosis, motility, angiogenesis, and immune responses, p53, one of the most important tumor suppressors, is inactivated in about half of all malignancies, and S1P has been implicated in tumor angiogenesis (52, 53, 54).

It would also be intriguing to explore whether DEK exerts differential effects on hormones such as estrogen, progesterone, and androgen in benign versus malignant gynecological tumors, considering these hormones’ established roles in regulating DEK expression and activity through various signaling pathways (55). For instance, studies show that estrogen can directly bind to specific receptors in the nucleus, leading to enhanced transcription of the DEK gene (55). This investigation would be particularly compelling given the key roles of these hormones in uterine myomas. For example, progesterone receptor isoform B mRNA levels in uterine myoma tissue have been found to correlate with tumor number and correlate inversely with intermenstrual bleeding and dysmenorrhea intensity, suggesting a role in uterine myoma growth and symptom attenuation (4). Furthermore, future research could investigate whether DEK expression varies throughout the menstrual cycle and whether these fluctuations are linked to hormonal changes. Studying postmenopausal tissues may also provide valuable insights into DEK’s expression patterns in the absence of hormonal cycles, helping to determine its stability or variability when hormonal influence is no longer present. Furthermore, an intriguing extension of this research would be to examine whether DEK expression levels are similar in the uterine cervix and the myometrium, particularly in the context of cervical cancer. By comparing DEK levels in cancerous cervical tissue to those in myometrial tissue that is unaffected by the tumor, we could gain valuable insights into DEK’s role in tumor biology and its interactions within the local microenvironment. This examination could help clarify whether DEK plays a specific role in the tumorigenic processes of cervical tissues or if its effects are more broadly applicable across different uterine tissues.

In conclusion, the significant upregulation of DEK in cervical cancer tissues, intermediate expression in uterine myomas, and negligible levels in normal uterine tissues highlight DEK’s potential role in tumor development and progression. These findings underscore the need for further research to elucidate the precise mechanisms by which DEK contributes to gynecological tumorigenesis and to explore its potential as a biomarker for early screening, thereby advancing the early detection and management of gynecological tumors and addressing a crucial need in women’s health.

## Declaration of generative AI and AI-assisted technologies in the writing process

During the preparation of this work the authors used ChatGPT in order to improve the readability and language of the manuscript. After using this tool, the authors reviewed and edited the content as needed and take full responsibility for the content of the publication.

## CRediT Authorship Contribution Statement

- Amelia Janiak: Conceptualization, Methodology, Formal Analysis, Investigation, Project Administration, Visualization, Writing - original draft
- Xinyue Liu: Validation
- Renata Koviazina: Writing - review & editing
- PengCheng Tan: Methodology, Investigation, Supervision, Validation, Writing – review & editing
- Ferdinand Kappes: Conceptualization, Methodology, Supervision, Writing - review & editing
- Fangrong Sheng: Resources, Writing - review & editing
- Felice Petraglia: Writing - review & editing
- Chiara Donati: Writing - review & editing
- Anastasia Tsigkou: Conceptualization, Methodology, Supervision, Project Administration, Funding Acquisition, Writing – review & editing

